## Supplemental Figure for "Dynamic metabolome profiling uncovers potential TOR signaling genes"

### Supplementary figure 1

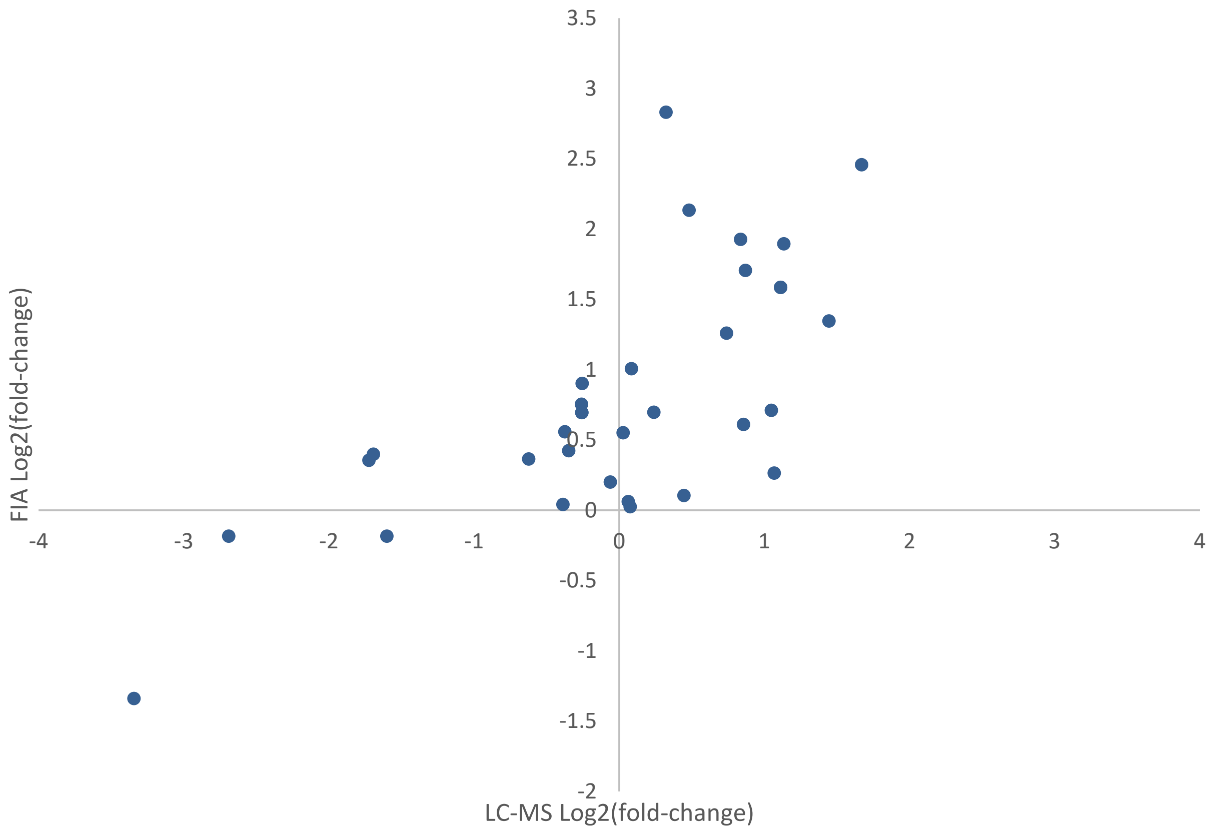


**Supplementary figure 1 – Comparison of effects of rapamycin treatment on the metabolome as measured by flow injection and LC-MS.** The average log2 transformed fold-changes for metabolites in wild-type yeast treated with 400 ng/mL rapamycin are compared between flow-injection and LC-MS measurements. Points indicate the average log2 fold change for three biological replicates for a metabolite that was measured using both analytical systems.

### Supplementary figure 2

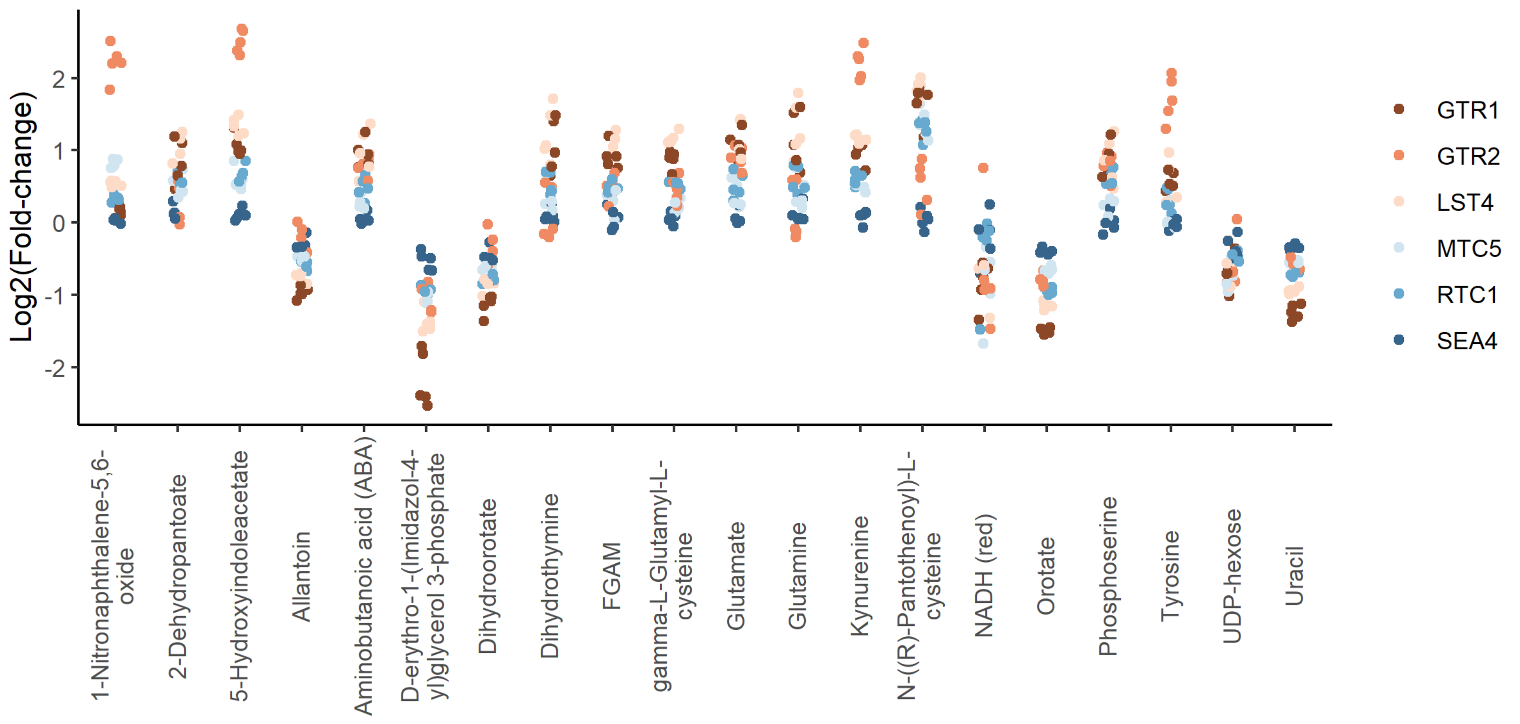


**Supplementary figure 2 – Positive regulators of TORC1 signaling show similar metabolite changes.** The Log2 transformed fold changes for each indicated mutant compared to the wild-type control are shown across the metabolites that demonstrated the largest changes in metabolite levels across the group. Each point indicates the fold change at one time point for one mutant.

### Supplementary figure 3

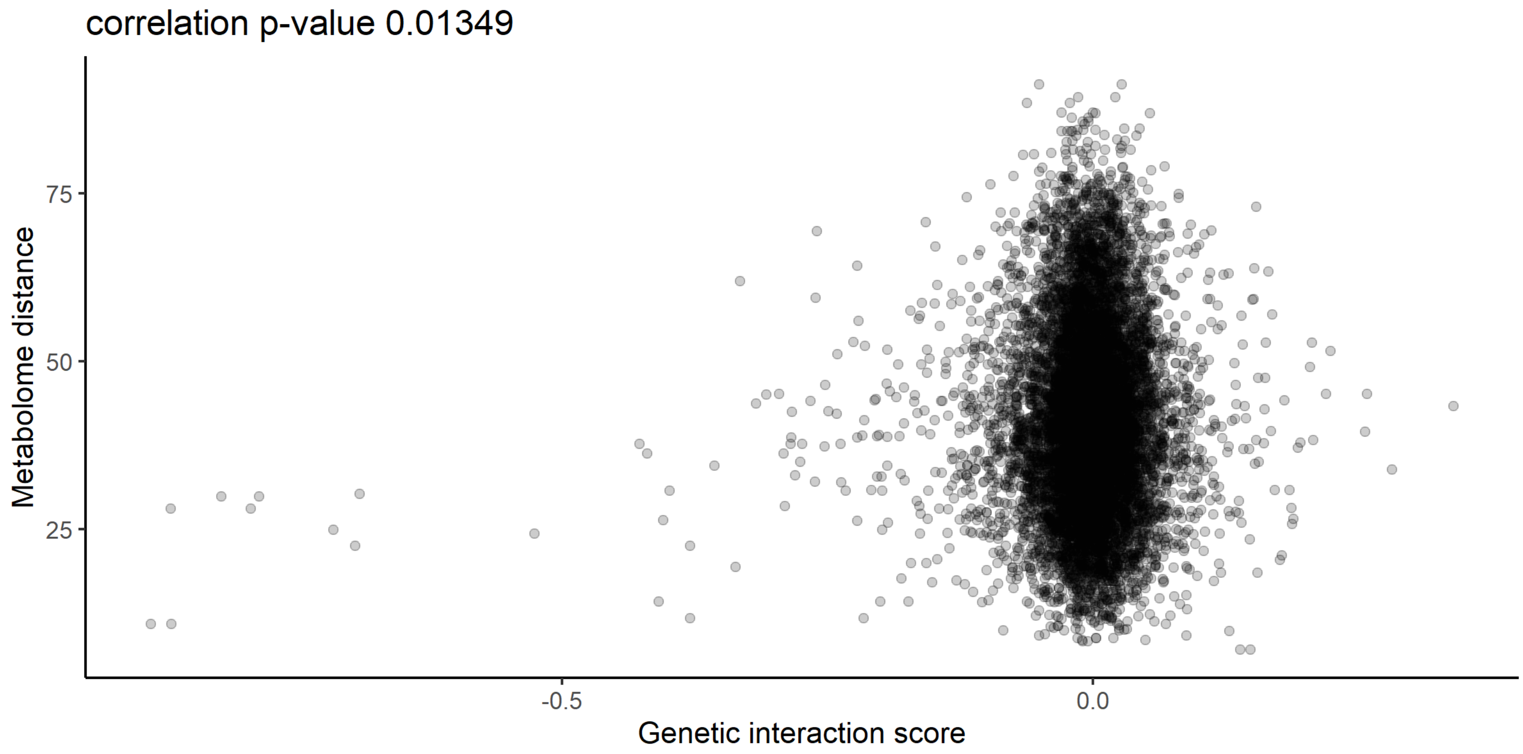


**Supplementary figure 3 – Relationship between metabolome distance and genetic interaction score.** The relationship between the metabolome distance as represented in Figure 2 between genes after 60 minutes of rapamycin treatment, and the genetic interaction score as defined in Costanzo *et al*[4].

### Supplementary figure 4

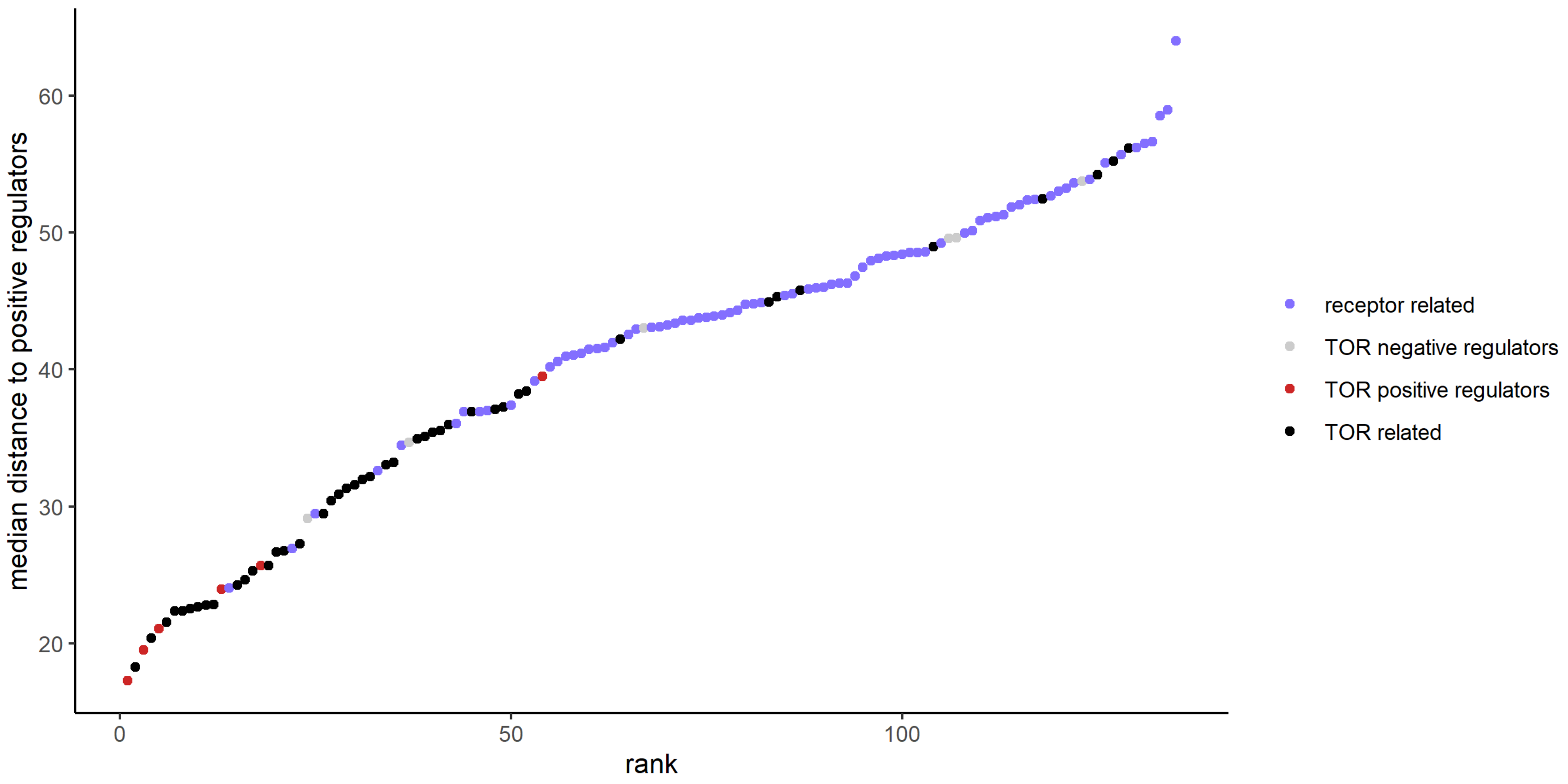


**Supplementary figure 4 – The metabolome distances between positive and negative regulators of TOR signaling.** The metabolome distance as represented in Figure 2 between genes after 30 minutes of rapamycin treatment and non-self, positive regulators of TOR signaling is shown in ascending order. Positive and negative regulators of TOR signaling are indicated by colour.

### Supplementary figure 5

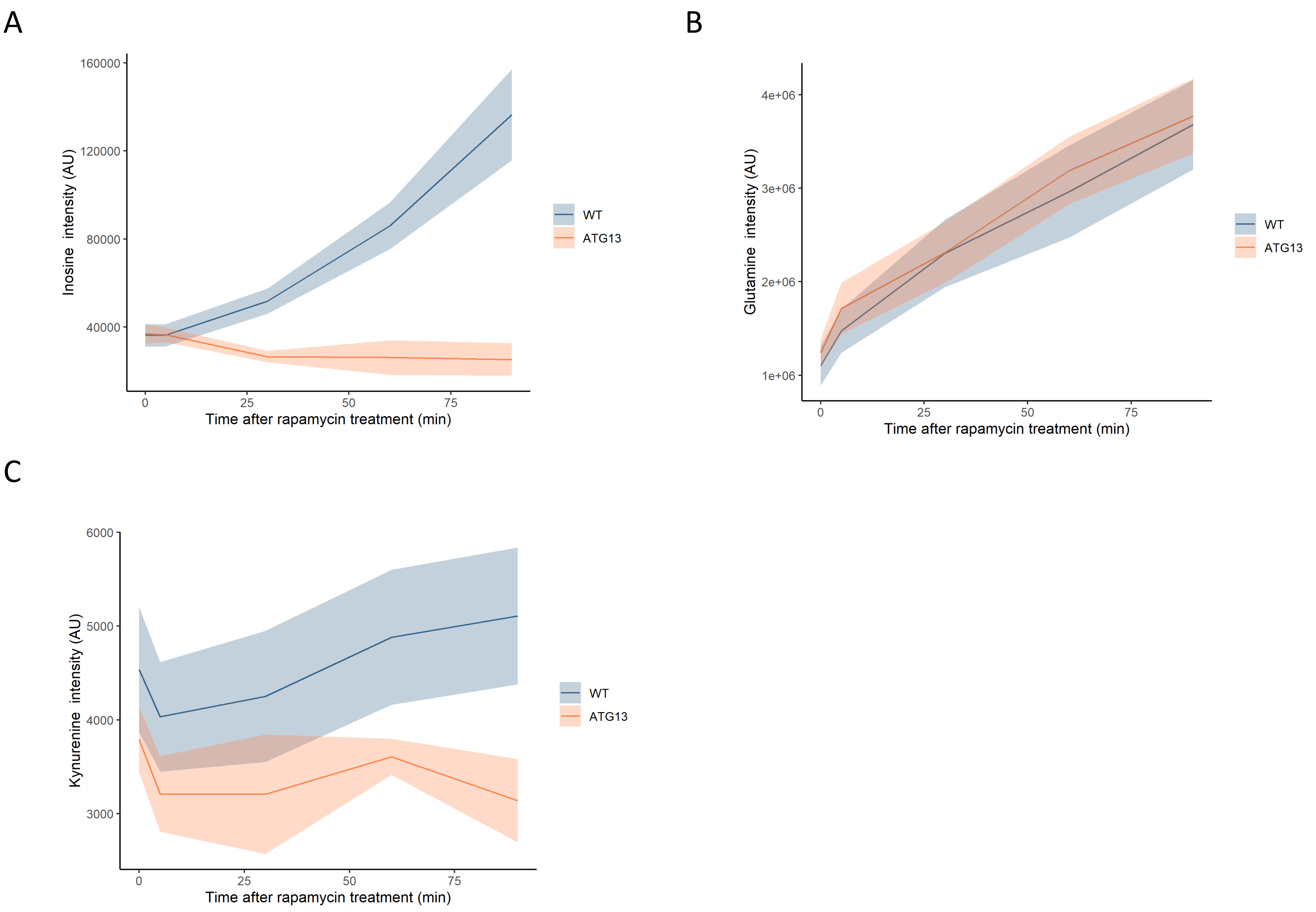


**Supplementary figure 5 – The effect of rapamycin on the levels of metabolites in wild-type and *atg13* yeast.** The normalized ion intensities for inosine (A), glutamine (B) and kynurenine (C) in wild-type and *atg13* yeast are shown after the indicated duration of rapamycin treatment. The central lines indicate the median value across 4 biological replicates. Shaded areas indicate the standard deviation across replicates.

Supplementary figure 6

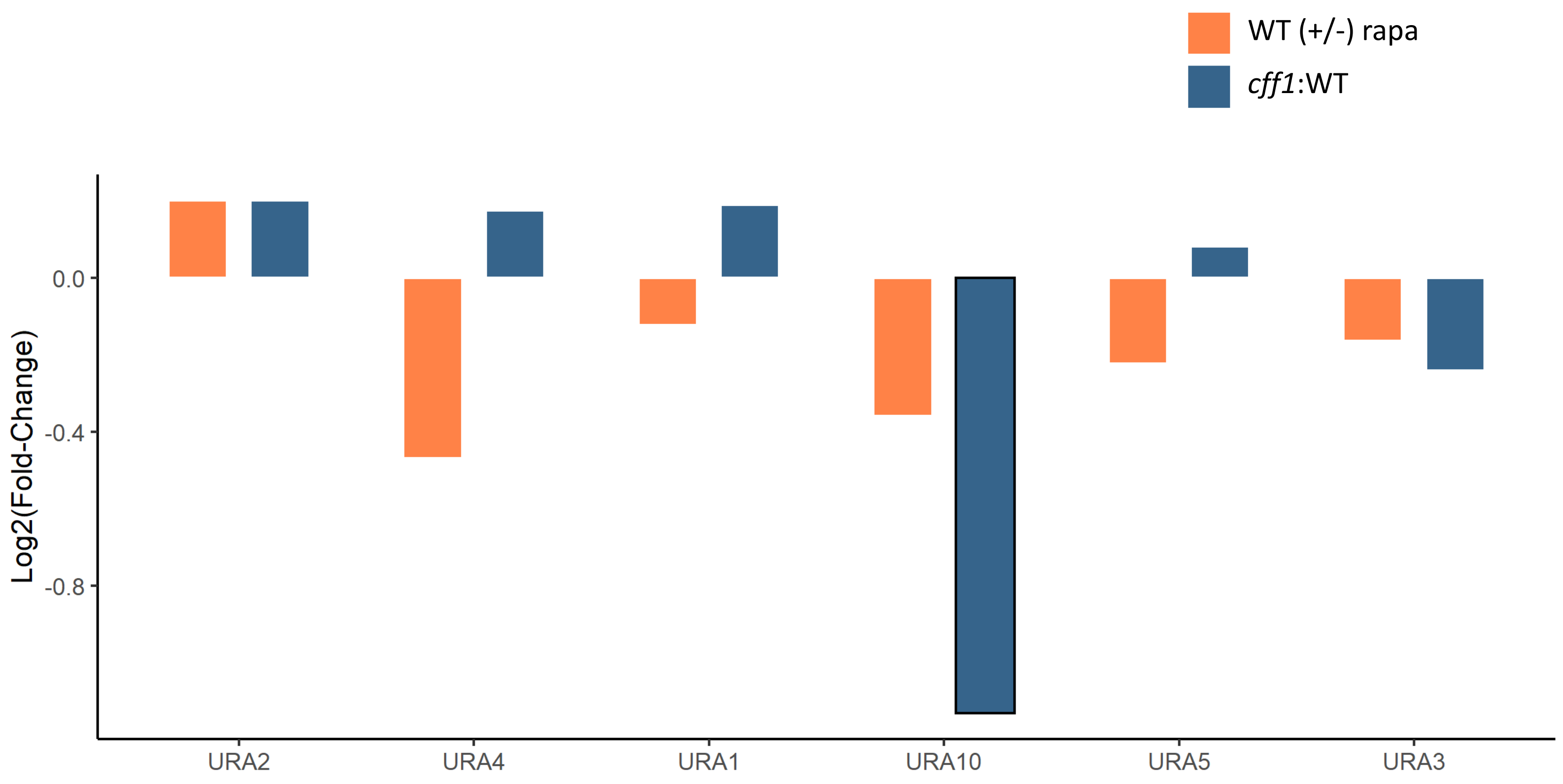


**Supplementary figure 6 – The effect of rapamycin and *CFF1* deletion on the levels of pyrimidine biosynthetic enzymes.** The average Log2 fold change for protein levels for the indicated comparison groups are shown (n = 3 biological replicates). Changes with a p-value of less than 0.05 (two sided Students T-test) are outlined in black.

### Supplementary figure 7

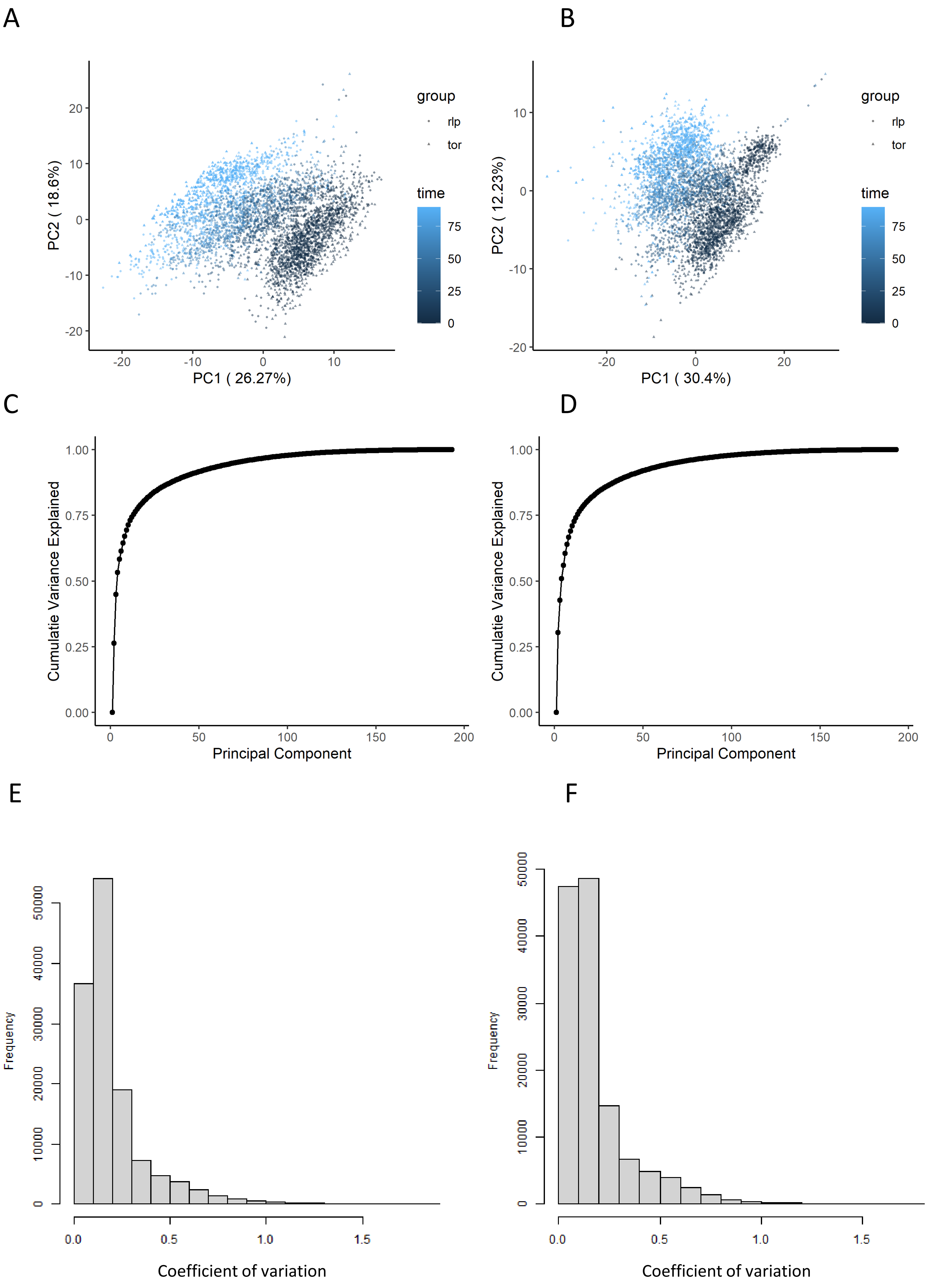


**Supplementary figure 7 – The effect of normalization of flow-injection metabolomics data.** AB) Principal components analysis representation of the variance within the dataset before (A) and after (B) the normalization as outlined in the methods section. The points are coloured according to the time after rapamycin treatment. CD) The cumulative variance explained by principal component analysis is shown before (C) and after (D) normalization. EF) The distribution of coefficients of variation for each ion and set of biological replicates is shown before (E) and after (F) normalization.

### Supplementary figure 8

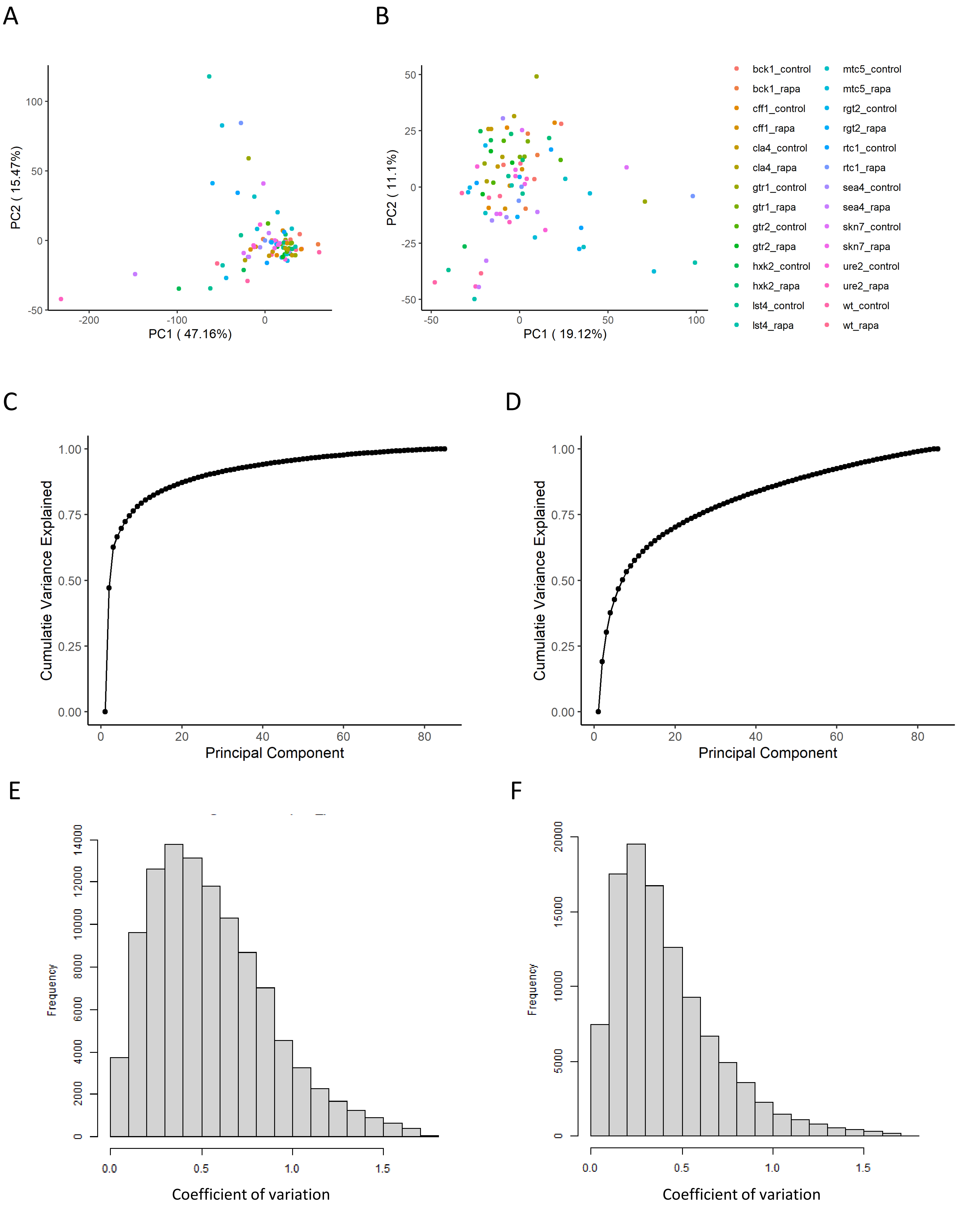


**Supplementary figure 8 – The effect of normalization of proteomics data.** AB) Principal component analysis representation of the variance within the dataset before (A) and after (B) the normalization as outlined in the methods section. The points are coloured according to the perturbation applied for the sample. CD) The cumulative variance explained by principal component analysis is shown before (C) and after (D) normalization. EF) The distribution of coefficients of variation for each ion and set of biological replicates is shown before (E) and after (F) normalization.
